# Oxidative phase transition of heat shock factor-1

**DOI:** 10.1101/2021.12.02.471034

**Authors:** Soichiro Kawagoe, Motonori Matsusaki, Koichiro Ishimori, Tomohide Saio

## Abstract

Heat shock factor 1 (Hsf1) was found as a central upregulator of molecular chaperones in stress adaptation, but it has recently been rediscovered as a major component of persistent nuclear stress bodies (nSBs). When the persistently stressed cells undergo apoptosis, the phase transition of nSBs from fluid to gel-like states is proposed to be an important event in switching the cell fate from survival to death. Nonetheless, how the phase separation and transition of nSBs are driven remain unanswered. In this study, we discovered that Hsf1 formed liquid-liquid phase separation droplets *in vitro*, causing the assembly of Hsf1 to drive nSBs formation. Under oxidative conditions, disulfide-bonded and oligomerized Hsf1 formed gel-like and more condensed droplets, confirmed through fluorescence recovery, refractive index imaging, and light scattering. Then, on the basis of our results, we proposed that Hsf1 undergoes oxidative phase transition by sensing redox conditions potentially to drive the cell fate decision by nSBs.

## Introduction

Organisms maintain homeostasis through both cell death induction and stress adaptation mechanisms, depending on environmental stress conditions^1^. Heat shock factor 1 (Hsf1) plays a central role in stress adaptation by upregulating molecular chaperones in mammalian cells^2^. Hence, an activation model of Hsf1 was proposed that Hsf1 existed as a monomer under normal conditions, but in response to stress, forms a trimer^3^. This Hsf1 trimer can then bind to the DNA sequence called the heat shock element as a transcription factor, which is located in the promoter region upstream of chaperone genes, thereby promoting the chaperone expression^4–6^. Chaperones are responsible for the refolding and anti-aggregation of misfolded proteins in the cytoplasm^7^.

While Hsf1 acts as a transcription factor in the nucleus during stress, it also constitutes nuclear stress bodies (nSBs)^8–10^ by binding to satellite III DNA repetitive sequences, which encode non-coding RNA. Microscopic observation of the cell showed that nSBs are transient membrane-less organelles that are formed under proteotoxic stress conditions as foci in nucleus, and resolve as the cell recovers from the stress^4,11,12^. Moreover, based on the reversibility of nSBs, the formation and resolution of nSBs is proposed to be controlled by liquid-liquid phase separation (LLPS)^13,14^. It has been shown that persistent stress induced the phase transition of nSB to a gel-like state, which is proposed to trigger apoptosis^13^. However, the mechanism of the formation and phase transition of nSBs in response to stress remains to be elucidated.

One of the potent regulators of nSBs is Hsf1. It has been shown that the cells that express Hsf1 molecules lacking the leucine-zipper domain (Fig. 1a), which is responsible for oligomerization of Hsf1, cannot form nSBs under continuous stress^8^, suggesting that the oligomerization of Hsf1 controls the formation and resolution of nSBs. Moreover, Hsf1 possesses an intrinsically disordered region that is often involved in LLPS formation (Fig. 1a)^15–17^. Although these observations imply that the assembly of Hsf1 can be responsible for both phase separation and transition of nSBs, this hypothesis has not been fully verified. Furthermore, the mechanism of the phase transition remains to be elucidated, as there has been no direct observation of Hsf1 droplet formation and the phase transition, and has been limited information about the architecture of the droplets.

**Fig. 1.**
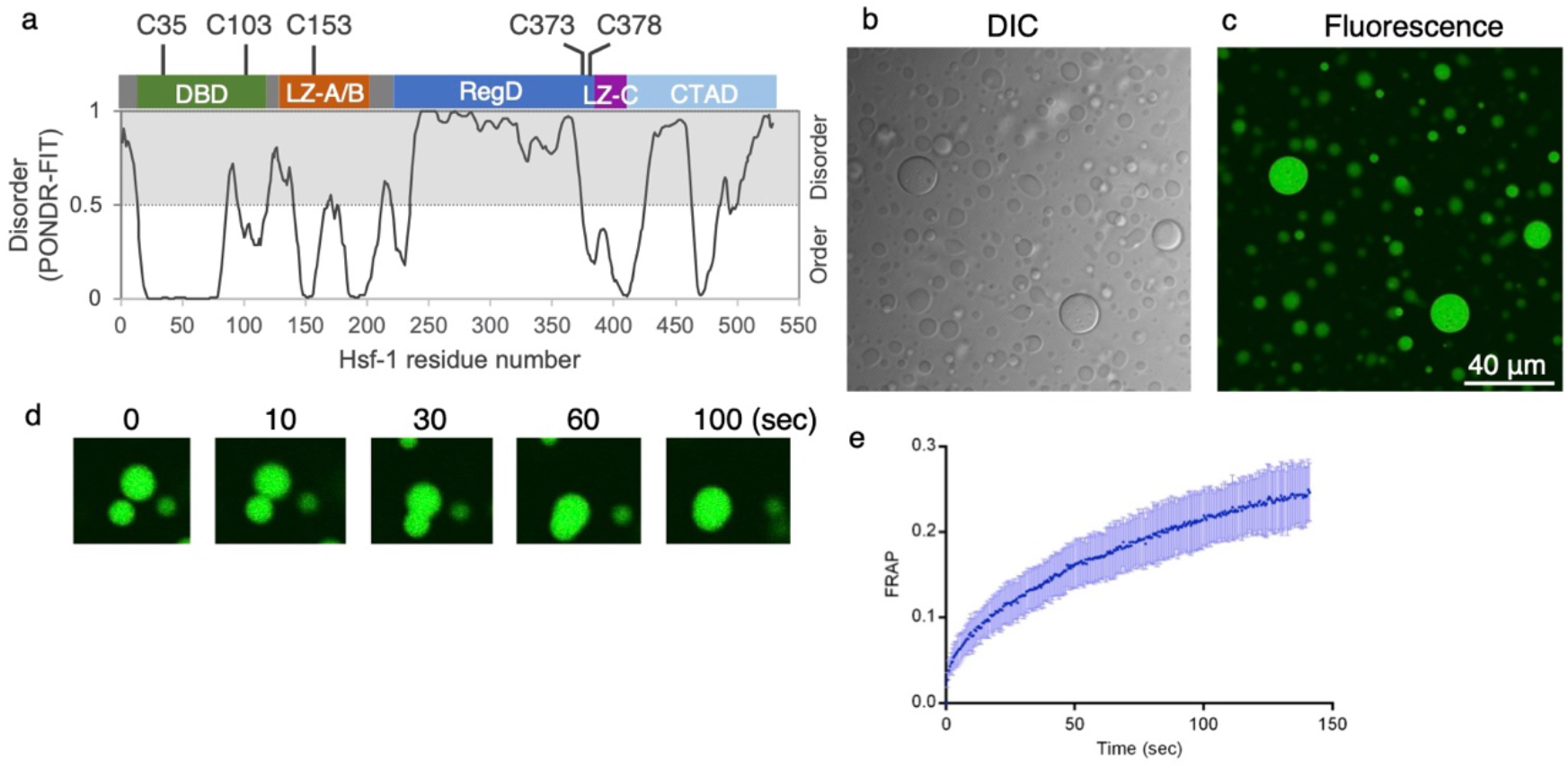
Hsf1 undergoes liquid-liquid phase separation *in vitro*. **a**, Domain organization, location of cysteine residues in Hsf1, and protein sequence and disorder prediction (PONDR) of Hsf1. The abbreviations used are: DBD, DNA-binding domain; LZ-A/B, leucine-zipper domain-A/B; RegD, regulatory domain; LZ-C, leucine-zipper domain; CTAD, C-terminal transactivation domain. **b, c**, Representative differential interference contrast and fluorescence image showing the droplets formed by Hsf1 (**b**) and Hsf1 with 0.1 eq Hsf1-GFP (**c**) in the presence of crowding agent and DTT. **d**, Time-lapse fluorescence microscopy showing fusion dynamics of Hsf1-GFP droplets. **e**, FRAP of a small fraction of the droplets is used to evaluate the internal mobility of Hsf1 droplets. The bleached event occurs at a time of 0 sec. The post-irradiation fluorescence intensity was normalized by the difference between the intensity before irradiation and the intensity of the first frame immediately after irradiation as 1 and recovery of the post-irradiation fluorescence intensity was analyzed. Data are plotted as mean ± s.d., with N = 7 independent experiments.

In this study, we aimed to elucidate Hsf1-driven droplet formations and regulatory mechanisms of the phase transition of Hsf1 droplets, which are the key event mediating the cell’s survival and death by the phase transition of nSBs. We first discovered that Hsf1 formed LLPS droplets *in vitro*, which proposed that the assembly of Hsf1 can drive phase separation of nSBs. Notably, since Hsf1 was oligomerized by intermolecular disulfide bonds^18^, the shape and internal mobility of Hsf1 droplets were then evaluated, focusing on the redox state. Conventional fluorescence recovery after photobleaching (FRAP) analysis confirmed that Hsf1 droplets were found to undergo phase transitions to gel-like droplet under oxidative conditions. Using holographic microscopy and size-exclusion chromatography and multi-angle light scattering (SEC-MALS) to focus on internal structure of Hsf1 droplet, we clearly showed that oxidative conditions also promoted the condensation of Hsf1 inside the droplet and favored the formation of higher-order Hsf1 oligomers. Thus, we clarified that the phase transition of Hsf1 droplets was redox-dependently driven by oligomerization, potentially to mediate the cell fate decision by nSBs.

## Results

### Hsf1 undergoes liquid-liquid phase separation

To verify whether Hsf1 formed liquid-like droplets, purified Hsf1 solutions under varying conditions were observed through microscopy. After adding crowding agents, the Hsf1 solution in the presence of a reducing agent; dithiothreitol (DTT), turned cloudy. We observed that the addition of a molecular crowder promoted droplet formation (Fig. 1b). Differential interference contrast microscopy images showed that a droplet having round globular shape was formed in the solution. Note that the enhanced LLPS-based droplet formation under crowding conditions as seen for Hsf1 was consistent with the previous reports on other low-complexity proteins showing that the LLPS was enhanced by the excluded volume effect of the molecular crowder^19–22^. The incorporation of Hsf1 in the droplet was further confirmed by confocal microscopy of droplets formed in the presence of 0.1 equivalent Hsf1-green fluorescent protein (GFP), showing GFP-derived fluorescence inside the droplets (Fig. 1c). Time-lapse imaging of the droplets using a confocal microscope showed that the droplets fused with each other in about one to two minutes (Fig. 1d), indicating the internal mobility of Hsf1 droplet to allow the rearrangement of Hsf1 molecules. The mobility of Hsf1 molecule in the droplet was also evaluated through FRAP^15^ analysis (Fig. 1e), in which a small portion of the droplet was bleached. Around 25% of the detected signal intensity of Hsf1-GFP droplets before bleaching was recovered within 140 sec after photobleaching (Fig. 1e). Our data therefore indicated that Hsf1 droplets were formed via LLPS, with enough mobility to fuse with each other under reductive conditions (Fig. 1d, e). These observations are consistent with the fluid character of nSBs found in cells^4^. Thus, Hsf1 is proposed to be a protein that drives nSBs formation in cells.

### Redox-dependent phase transition of Hsf1

Although it has previously been shown that the fluid-to-gel-like state transition of nSBs progresses in proportion to the duration and intensity of the stress^13^, the mechanism of transition has not been elucidated. Interestingly, it has been reported that reactive oxygen species (ROS) are produced in cells by exposure to stresses, such as heat and heavy metals^23–25^, implying a possible relationship between oxidative conditions and phase transitions of nSBs. Therefore, to evaluate the effect of oxidative conditions on the Hsf1 droplet, we investigated the Hsf1 droplet *in vitro* under reductive and oxidative conditions. We observed that Hsf1 droplets in the presence of the oxidizing agent (H_2_O_2_) or in the absence of the reducing agent were in distorted gel-like shapes, in which the small droplets were assembled but not fused into one droplet (Fig. 2a). Such distorted shapes are characteristic of LLPS droplets altered from mobile to immobile states^26^, implying that the Hsf1 droplet undergoes phase transition under oxidative conditions.

**Fig. 2.**
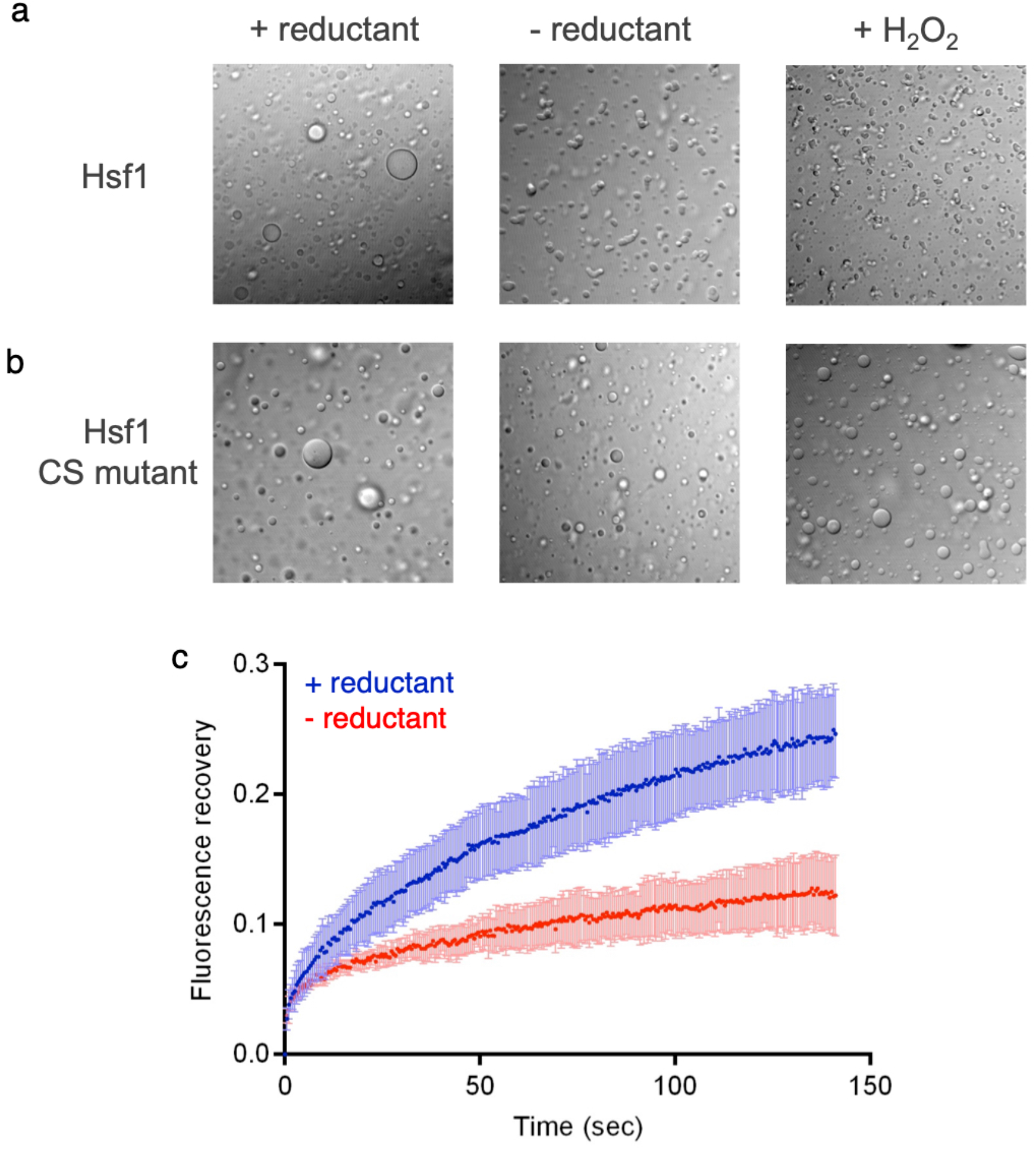
Redox-dependent phase transition of Hsf1. **a**,**b**, Differential interference contrast images of Hsf1 droplets (**a**) and Hsf1 CS mutants droplets (**b**) in the presence of crowding agent. Left panels are in the presence of DTT (+ reductant), center panels are in the absence of redox agents (− reductant), and right panels are in the presence of H_2_O_2_. **c**, FRAP data shows the decrease of mobility of Hsf1 droplet at the oxidative condition. The bleached event occurs at a time of 0 sec. Data are plotted as mean ± s.d., with N = 7 independent experiments.

Since the oxidative conditions were found to drive the external morphological changes of LLPS droplets, we next investigated their internal structure. To evaluate the internal mobility of Hsf1 droplets under oxidative conditions, FRAP experiments were conducted. After bleaching small portions of the droplet to evaluate the mobility of Hsf1 molecules in the droplets over time, the degree of fluorescence recovery was found to be smaller under oxidative conditions than under reductive conditions, showing the decreased internal mobility of Hsf1 droplet in the oxidative condition (Fig. 2c). Collectively, shape transformation and internal mobility evaluations showed that Hsf1 undergoes phase transition to the gel-like droplet in the oxidative condition.

It is known that Hsf1 directly senses heat stress^18,27,28^, but the mechanism for sensing redox environment is not understood. A redox environment can be sensed via Cys residues that can form inter- or intramolecular disulfide bonds. Hsf1 has five cysteine residues (Fig. 1a), and it has been suggested that heat stress induces intracellular Hsf1 oligomerization by intermolecular disulfide bond formation^18,27^. Thus, to evaluate the effect of disulfide bond formations on the properties of Hsf1 droplets, the Hsf1 CS mutant that lacks all five Cys residues was prepared. As a result, shapes of the Hsf1 CS mutant droplet remained spherical, even under oxidative conditions (Fig. 2b), suggesting Hsf1 CS mutant lost the ability to sense redox environments.

### Oxidative environments promote the accumulation of highly-condensed structures inside the Hsf1 droplet

Since it was shown that the Hsf1 droplet undergoes a redox-dependent phase transition, we further analyzed the internal structure of the Hsf1 droplet to uncover the mechanism of the phase transition. We attempted to visualize the internal structure of droplets under redox conditions using a holotomography imaging that visualizes the object based on the refractive index (RI)^29^. We observed by holotomography microscope that Hsf1 droplets were formed after adding crowding agent. Afterward, the 3D shape of the Hsf1 droplet based on RI was visualized. As seen by confocal microscopy observation (Fig. 2a), the shape of Hsf1 droplets was round and globular under reductive conditions and distorted under oxidative conditions (Fig. 3a). Quantitative evaluations of RI can also be used to evaluate the distribution of the Hsf1 molecule inside the droplet. RI mapping showed that RI values were not uniform in the droplet, and regions with higher RI values were sparsely scattered in the droplets, indicating that Hsf1 molecules in the droplet were not uniformly distributed (Fig. 3b).

**Fig. 3.**
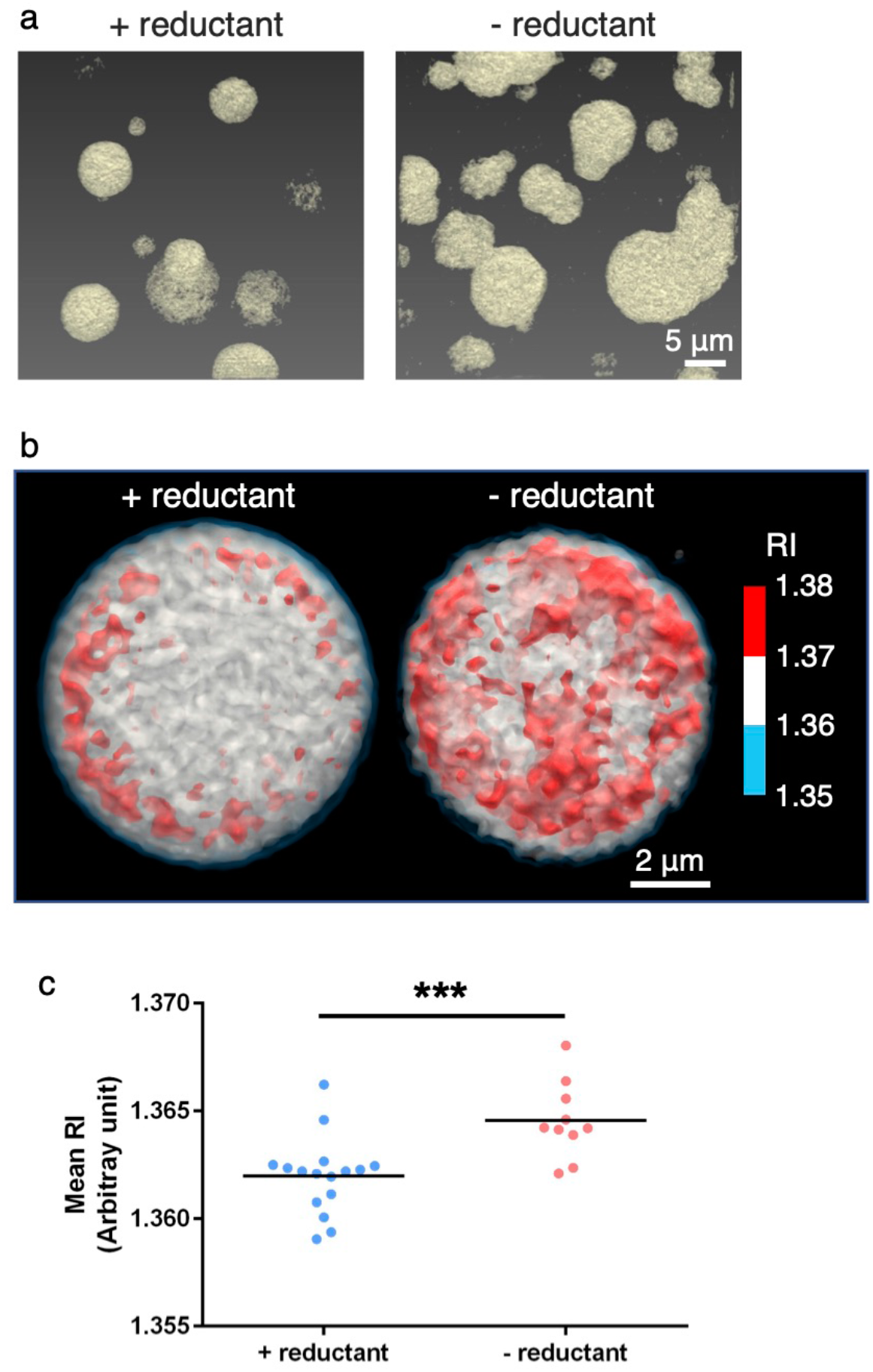
Observation of internal state of Hsf1 droplet using holographic microscope. **a**, 3D RI images of Hsf1 droplet. Left and right panel refer to the RI images of Hsf1 droplet with or without DTT (+,− reductant), respectively. The areas with RI above 1.353 are colored yellow. **b**, Quantitative analysis of tomographic images, measuring the mean RI of the Hsf1 droplet in the presence of DTT (+ reductant) and in the absence of DTT (− reductant). **c**, A scatter plot with N = 16 (+ reductant) and 10 (− reductant). Each plot shows the mean RI of the average of the individual droplets. ***, *p* < 0.001, statistical significance between mean RI of + reductant and − reductant.

Regions with higher RI values are expected to have higher concentrations of the molecule, suggesting the existence of “core particles” in the droplet. Interestingly, comparison between RI mapping of droplets under reduced and oxidized conditions highlighted that the droplets in the oxidative condition had more contents of the core particle than those in reductive conditions (Fig. 3b). Therefore, to quantitatively evaluate the RI inside Hsf1 droplets, several droplets of 3−8 μm were selected, after which the mean RI of each droplet was calculated to make a scatter plot. The scatter plot showed that droplets in the oxidative condition had significantly higher mean RI values than those subjected to reductive conditions (Fig. 3c), indicating that droplets in the oxidative condition had higher concentrations of the protein on average. Likewise, on the basis of FRAP data showing a lesser mobility of the Hsf1 molecule in the droplet formed during oxidative conditions (Fig. 2c), the higher density and more content of core particles in the oxidative droplet can be key features in understanding the mechanism of phase transitions. Therefore, our data suggest that the tighter packing of Hsf1 molecules leads to gel-like droplet properties.

### The promotion of Hsf1 oligomer assemblies through disulfide bonds

Holotomography showed that the Hsf1 droplet in the oxidative condition was characterized by a higher density, consisting of more core particle contents. To unveil the mechanism of the enhanced formation of the core particles in the oxidative condition, the oligomeric state of Hsf1 in the reduced and oxidative conditions were evaluated by SEC-MALS (Fig. 4a, b, c). To evaluate the molecular assembly of Hsf1, SEC-MALS experiment was performed for Hsf1 in the soluble oligomeric state. In the presence of DTT, the absorbance peak at 280 nm from the gel filtration chromatography was highly symmetric. We also observed that the median molar mass of Hsf1 at an injection concentration of 50 μM was 379 kDa, corresponding to the 7-mer (Fig. 4a, d). Alternatively, in the absence of DTT, the symmetry of the absorbance peak was broken and the median molar mass increased to 540 kDa, corresponding to 9-mer (Fig. 4b, d). Furthermore, more components with larger molar mass exist in the absence of DTT, which can form higher-order oligomers up to 16-mer. Hsf1 CS mutants without DTT did not form such higher-order oligomers and the median molar mass was 308 kDa, corresponding to 5-mer (Fig. 4c, d). Hence, these results propose that Hsf1 responds to oxidative conditions at the molecular level and forms a higher-order oligomer via Cys residues. Consequently, RI of the core particle observed through holotomography imaging can be interpreted to reflect the assembly state of Hsf1 oligomers in the droplet.

**Fig. 4.**
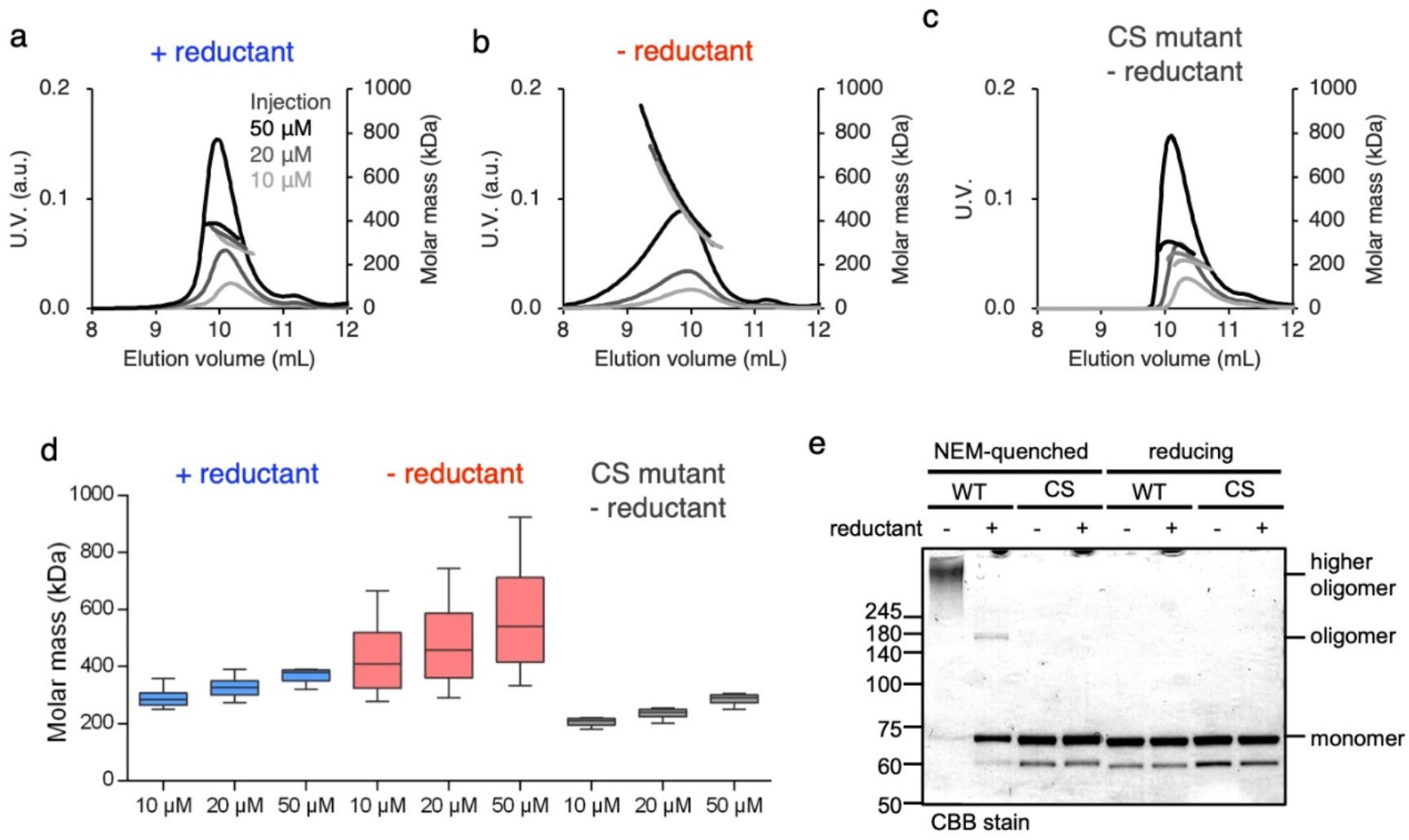
Redox-dependent Hsf1 oligomerization. **a**,**b**,**c**, SEC-MALS of Hsf1 with DTT (+ reductant) (**a**), without DTT (− reductant) (**b**), and Hsf1 CS mutant without DTT (CS mutant – reductant) (**c**) injected at varying concentrations. **d**, A box and whisker plot of Hsf1 molecular mass at various conditions. **e**, Redox states of HSF1 under the reductive condition. HSF1 was incubated in the absence or presence of DTT. The samples were quenched by a large excess of NEM (NEM quenched) or SDS-sample buffer containing a reductant (reducing), and then the samples were separated by SDS-PAGE. Red and ox mean the Hsf1 with or without DTT, respectively.

To verify that this higher-order oligomer was formed through intermolecular disulfide bonds between Hsf1 molecules, SDS-PAGE was performed. Results detected the Hsf1 WT and CS mutant as monomers in reducing SDS-PAGE. However, in N-ethylmaleimide (NEM) quenched non-reducing SDS-PAGE, higher-order oligomers were detected only in Hsf1 WT prepared under oxidative conditions (Fig. 4e). Thus, this result confirms that the higher-order oligomer formation of Hsf1 was driven by intermolecular disulfide bonds.

## Discussion

The formation of nSBs is an important event that regulates cell fate. However, the complexity of its composition has concealed the detailed mechanism of generating and regulating nSBs. This study therefore focused on Hsf1, a major component of nSBs. We observed that the purified Hsf1 molecules formed LLPS droplets in the crowded environment (Fig. 1b, c), suggesting that LLPS of Hsf1 contributed to the formation of nSBs. Interestingly, microscopic observations and FRAP measurements unveiled that the fluidity of the Hsf1 droplet was altered by redox conditions, causing droplets to undergo phase transition from liquid to gel-like states under oxidative conditions (Fig. 2a, c). Hence, the redox-dependent phase transition of Hsf1 suggested a cause of the phase transition of nSBs under continuous stress^13^. The fact that ROS molecules were produced in cells exposed to stresses also supported this ideas^23–25^. Thus, given the fact that the phase transition of nSBs leads to apoptosis^13^, our data propose that phase transition of Hsf1 is a key factor responding to continuous stress for regulating apoptosis.

Phase transition mechanisms of Hsf1 droplets have been addressed using holotomography imaging to identify the architecture of these droplets (Fig. 3b). Mapping of RI values showed that Hsf1 molecules were inhomogeneously distributed. Core particles with higher density were also formed in the droplet. Furthermore, data showed that oxidative conditions promoted the formation of core particles, thereby leading to more condensed droplets (Fig. 3b, c). Moreover, SEC-MALS data show that Hsf1 molecules formed higher-order oligomers, consisting of up to ~16 Hsf1 molecules under oxidative conditions (Fig. 4b, d). The oligomers observed in the SEC-MALS experiment are expected to have intermolecular interactions with each other when they drive LLPS droplet formation in the presence of the molecular crowder. Larger oligomers, consisting of a larger number of Hsf1 molecules are also expected to have a higher affinity with other oligomers, which results in the promoted formation of core particles (Fig. 5).

**Fig. 5.**
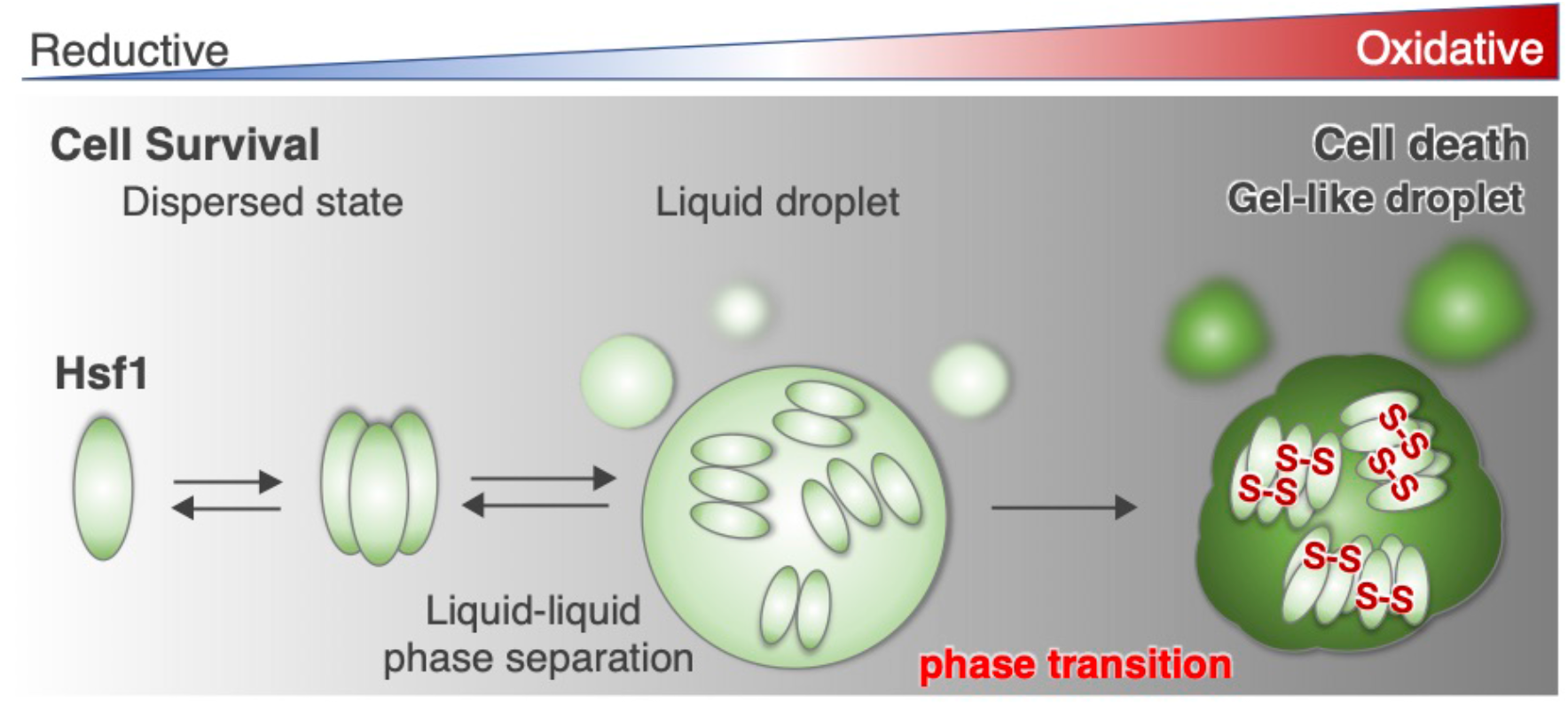
The model of phase transition of Hsf1 droplet. Hsf1 forms LLPS droplets upon accumulation and undergoes phase transition to gel-like droplet in a redox-dependent manner. Phase transition of Hsf1 is driven by the formation of higher-order oligomers via disulfide bonds between Hsf1 molecules. The oxidative phase transition of Hsf1 may lead to cell death.

As described above, our data show that Hsf1 exists in larger oligomers than that modelled in the previous studies^2,3,30^. It has been reported that Hsf1 existed as a monomer in the cytoplasm under normal conditions, and stress conditions induced the trimerization of Hsf1, leading to translocation into the nucleus^2,30–32^. Alternatively, several previous studies on electrophoresis and gel filtration column chromatography indicated that Hsf1 formed a larger oligomer than the trimer^18,33,34^. In this study, quantitative analysis using SEC-MALS showed that Hsf1 formed larger oligomers than trimers, with ~7-mer being formed in the reduced state, and ~9-mer being formed in the oxidized state as a median molar mass. The formation of the larger oligomer can be important for LLPS in the stress response.

In oxidized conditions, the formation of disulfide bonds (Fig. 4e) contributes to the assembly of Hsf1 molecules, which is further corroborated by using the Hsf1 CS mutant. Interestingly, microscopic observations also showed that Hsf1 CS mutant droplets had a similar morphology to reduced Hsf1 droplets, even in an oxidative environment in the presence of H_2_O_2_ (Fig. 2b). This result suggested that Hsf1 CS mutant droplets cannot undergo a phase transition. Hsf1 has five Cys residues, and both intramolecular and intermolecular disulfide bond formations were expected. Given the fact that the disulfide bond formation makes the molecule more compact^35^, the disulfide bond formation of Hsf1 can make the protein more compact, which contributes to the formation of more condensed core particles (Fig. 2a, 3a, b). Collectively, we propose a mechanism of phase transition, where the synergistic effect of the packing of the Hsf1 oligomer by disulfide bonds and an increase in intermolecular interactions due to the formation of higher-order oligomers, promoted the formation of core particles, which eventually resulted in a higher density and lower fluidity inside the droplet (Fig. 5). Such a mechanism of redox condition-dependent phase transition of Hsf1 droplets can explain the nSBs phase transition under continuous stress and potentially mediates the cell fate decision by nSBs.

In addition to Hsf1, a few components of nSBs, including RNA polymerase II and a disordered region of the bromodomain-containing protein 4 (BRD4), have been reported to undergo phase separation *in vitro*^22,36^, suggesting that nSB is a membrane-less organelle composed of multiple LLPS-potent proteins. Both RNA polymerase II and BRD4 are involved in the transcription of the satellite III RNA^37,38^ that works as a scaffold to assemble the other nSB components to regulate the intracellular splicing of genes associated with multiple functions, including DNA/RNA metabolism, biosynthesis, stress response, and cell cycle^39,40^. Therefore, the LLPS and redox-dependent phase transition of Hsf1 imply the contribution of redox condition to the regulation of the Sat III RNA transcription through nSBs.

## Materials and methods

### Expression and purification of protein samples

The *human* Hsf1 and Hsf1-GFP expression constructs were cloned into a pET vector (Novagen). Hsf1 CS mutant (C35S/C103S/C153S/C373S/C378S) was then constructed through site-directed mutagenesis using PrimeSTAR Mutagenesis Basal Kit (Takara Bio). All the protein samples were expressed in BL21(DE3) and purified using Ni-NTA Sepharose column (QIAGEN) and Superdex 200 16/600 column (GE Healthcare).

### Confocal microscopy

Fluorescence images of the Hsf1 droplets were obtained using a confocal microscope (FV1200, Olympus).

### Fluorescence recovery after photobleaching

Fluorescence recovery after photobleaching (FRAP) experiments was conducted on *in vitro* droplets formed by Hsf1, which was mixed with Hsf1-GFP, using the 473 nm laser line of a confocal microscope (FV1200, Olympus). For each droplet, either the whole droplet or a spot was bleached for 2–10 s, after which post-bleach time-lapse images were collected. The resulting images were analyzed as follows: A 5-µm diameter region of interest (ROI) was placed on the bleached whole droplet or a bleached spot. Fluorescence intensity of the ROI was then calculated using FV10-ASW (Olympus). Afterward, the post-irradiation fluorescence intensity was normalized, using the difference between the intensity before irradiation and the intensity of the first frame immediately after irradiation as one. Finally, the recovery of the post-irradiation fluorescence intensity was analyzed using Prism 5 (GraphPad Software).

### SEC-MALS experiments

Size-exclusion chromatography with multi-angle light scattering (SEC-MALS) was measured using DAWN HELEOS8+ (Wyatt Technology Corporation), a high-performance liquid chromatography pump LC-20AD (Shimadzu), a refractive index detector RID-20A (Shimadzu), and a UV-vis detector SPD-20A (Shimadzu) that was located downstream of the Shimadzu liquid chromatography system. The differential refractive index (Shimadzu Corporation) downstream of MALS was then used to obtain protein concentrations. The data were analyzed with ASTRA version 7.0.1 (Wyatt Technology Corporation).

### Holotomography imaging

Three-dimensional (3D) quantitative phase imaging and its correlative fluorescence live BM2 cell images were obtained using a commercial holotomography instrument (HT-2H, Tomocube), which is based on Mach–Zehnder interferometry, equipped with a digital micromirror device. The coherent monochromatic laser (λ = 532 nm) was divided into two paths, a reference, and a sample beam, respectively, using a 2 × 2 single-mode fiber coupler. Afterward, the visualization of 3D RI maps was performed using a commercial software (TomoStudio, Tomocube). The protein volume condensates were subsequently calculated by considering the physical size of individual voxels. The mean RI of each droplet was also calculated using Prism 5 (GraphPad Software).

### Gel-based analysis of HSF1 redox states

HSF1 was incubated with a reaction buffer for 1 h. To prepare NEM quenched samples, a 12 µL aliquot was taken from the reaction mix, after which 48 µL of NEM (Cat. No. 15512-24; Nacalai Tesque, Kyoto, Japan) was added to quench the reaction. Additionally, 60 µL of Laemmli 4 × sodium dodecyl sulfate (SDS)-sample buffer^41^ was added to these samples. Then, they were separated by SDS-PAGE. To prepare reducing samples, a 12 µL sample aliquot, 48 µL distilled water, and 60 µL SDS-sample buffer, containing β-mercaptoethanol (Cat. No. 15512-24; Nacalai Tesque) were mixed to reduce and denature the samples.

### Disorder predictions

Intrinsically disordered Hsf1 regions were predicted, using the “VSL2” algorithm of “Predictor of Natural Disordered Regions” (PONDR, http://www.pondr.com/).

## Acknowledgments

T.S. thanks Dr. Eiichiro Mori for fruitful discussion in the conceptualization of the study. This work was supported by grants from JSPS KAKENHI (JP20J20761 to S.K., 20K15969 to M.M., 17H05657, 17H05867, 18H05229, and 19K06504 to T.S., and 19H05769 to K.I.), MEXT Grant-in-Aid for Transformative Research Areas (B) (JP21H05094 to T.S.), JST FOREST Program (JPMJFR204W) to T.S.), and HIRAKU-Global Program, which is funded by MEXT’s “Strategic Professional Development Program for Young Researchers” to M.M. We also thank Takaki Ushifusa (Shinkouseiki. Co., Ltd.) for his help with holotomography imaging. This research in part used experimental resources provided by Fujii Memorial Institute of Medical Sciences, Institute of Advanced Medical Sciences, Tokushima University.

## Author contributions

S.K., M.M., and T.S. designed the research. S.K. and M.M. conducted the research. S.K. and M.M. analyzed the data. S.K., M.M., K.I., and T.S. wrote the paper with input from all other authors. T.S. conceptualized the study.

